# Activation of the dorsal septum increases alcohol consumption in male C57BL/6J mice

**DOI:** 10.1101/2022.01.15.476378

**Authors:** Harold L. Haun, Shannon L. D’Ambrosio, Thomas L. Kash

**Author notes:** **CORRESPONDING AUTHOR:** T. L. Kash, Bowles Center for Alcohol Studies, University of North Carolina at Chapel Hill, CB 7178 Thurston Bowles Building, 104 Manning Drive, Chapel Hill, NC 27599.

## Abstract

Binge drinking is a common pattern of excessive alcohol consumption associated with Alcohol Use Disorder (AUD) and unraveling the neurocircuitry that promotes this type of drinking is critical to the development of novel therapeutic interventions. The septal region was once a focal point of alcohol research yet has seen limited study over the last decade in relation to binge drinking. Numerous studies point to involvement of the dorsal septum (dSep) in excessive drinking and withdrawal, but few studies have manipulated this region in the context of binge drinking behavior. The present experiments were primarily designed to determine the effect of chemogenetic manipulation of the dSep on binge-like alcohol drinking in male and female C57BL/6J mice. Mice received bilateral infusion of AAVs harboring hM4Di, hM3Dq, or mCherry into the dSep and subjects were challenged with systemic administration of clozapine-*N*-oxide (CNO; 3 mg/kg) and vehicle (saline; 0.9%) in the context of binge-like alcohol consumption, locomotor activity, and sucrose drinking. CNO-mediated activation (hM3Dq) of the dSep resulted in a significant increase in binge-like alcohol consumption, locomotor activity, and sucrose intake in male mice. DSep activation promoted sucrose drinking in female mice, but alcohol intake and locomotor activity were unaffected. Conversely, silencing (hM4Di) of the dSep modestly decreased locomotor activity in males and did not influence alcohol or sucrose intake in either sex. Lastly, CNO was without effect in mCherry-expressing control groups. These data support a role for the dSep in promoting excessive, binge-like drinking behavior in a sex-dependent fashion and suggests a broad role for the region in the modulation of locomotor activity as well as general appetitive behavior such as sucrose drinking.

## 1. INTRODUCTION

The COVID-19 pandemic has had a palpable negative impact on mental health and incidences of binge drinking have seen a concomitant rise in the US [1,2]. Binge drinking is defined as the rapid consumption of alcohol such that blood alcohol concentrations (BAC) exceed the 0.08 g/dL legal limit of intoxication in a relatively short period of time [3–5]. This type of drinking is the most common pattern of excessive alcohol consumption and is associated with an increased risk for the development of an alcohol use disorder (AUD) [6-10]. Alcohol dependence and AUDs create a sizeable social and economic impact amounting to roughly $249 billion annually and is a leading cause of preventable mortality [11–13]. However, treatment strategies are severely lacking and a more thorough understanding of the neurobiological processes that govern excessive, uncontrolled alcohol drinking is necessary to meet this goal.

Preclinical models including the Drinking-in-the-Dark (DID) paradigm serve as a platform for pharmacological and circuit-level interrogation of systems that drive binge drinking, paving the way for the development of much needed novel pharmacotherapies [14–16]. Subcortical limbic structures such as the septal complex, hippocampus (HPC), amygdala (AMY), and nucleus accumbens (NAc) are of interest in promoting binge drinking behavior. In fact, researchers have demonstrated involvement of the HPC [17,18], AMY [19,20], and NAc [21,22] in excessive binge-like drinking behavior. However, no studies to date have manipulated the septal complex in the context of binge drinking and this is an intriguing avenue for further study given the role of this region in various alcohol related behaviors.

The septal region was one of the first structures identified by Olds and Milner to be responsive to electrical self-stimulation protocols, which suggested involvement in reward-related behavior [23]. Around the same time, the septum garnered much attention as lesion resulted in the phenomenon known as “septal rage” where rodents display highly aggressive behavior [24,25]. Much progress has been made since this time and the septum has been found to be an integral node in circuitry mediating spatial navigation and movement, reward processing, social behaviors, mood/affect, and a key component in generating theta rhythms and oscillations [26]. Indeed, patients diagnosed with an AUD display many behavioral disturbances consistent with septal function, such as movement disorders [27], reward-deficit biased behavior [28], increased aggressiveness and negative affective behavior [29,30], and altered theta [31]. Given these functions, it is likely that the septum plays a role in the various behaviors displayed in AUD and is a candidate for further study.

Interestingly, study of the septum in the context of AUD is somewhat limited. Human imaging studies found decreased septal volume in patients with an AUD or Wernicke-Korsakoff Syndrome [32,33]. Preclinical studies canvasing the effect of alcohol drinking on expression of the immediate early gene, c-fos, used this marker as a proxy for neuronal activity and found decreased expression following systemic alcohol challenge and voluntary consumption [34,35]. Further electrophysiological studies revealed that alcohol enhanced GABAergic activity to promote septal inhibition in rats with a history of drinking [36,37]. These findings suggest that the septum is responsive to acute alcohol challenge and that neuroadaptations occur with chronic consumption. Furthermore, c-fos activity is increased within the septum during withdrawal [38], suggesting engagement of the septal complex during a time when relapse-like behavior is observed. This general hypothesis is supported by work from the Aston-Jones lab that found increased septal cfos activity during cocaine-seeking behavior [39]. Furthermore, inactivation of the septum blocked contextual- and cue-induced reinstatement for cocaine [40], and attenuates motivated responding [41], supporting a role for the septum in relapse-like behavior.

While novel techniques within the preclinical research toolkit have advanced greatly in the last decade, studies targeting the septum have been largely absent from modern literature in the alcohol field. To fill this void, we tested the role of the dorsal septum (dSep) in the context of binge-like alcohol drinking behavior in male and female C57BL/6J mice using a chemogenetic approach. The primary objective of this study was to assess the role of the dSep on alcohol intake, however, the septal region also modulates locomotor activity and general reward related behavior. Thus, additional experiments were conducted to determine the contribution of the dSep to locomotor activity in an open field task and on sucrose consumption.

## 2. MATERIALS AND METHODS

### 2.1. Subjects

Male and female (N= 30/sex) C57BL/6J mice (Jackson Laboratories, Bar Harbor, ME) at 10 weeks of age were singly housed and tested in a temperature and humidity controlled AAALAC approved facility on a reverse 12-hr light/dark cycle. For all experiments, rodent chow (Isopro, RMH300) and water were provided *ad libitum* and mice were treated in accordance with both the NIH Guide for the Care and Use of Laboratory Animals and the Institutional Animal Care and Use Committee at UNC.

### 2.2. Surgical Procedures

Mice were anaesthetized with isoflurane (2%) and the AAV construct was bilaterally infused into the dSep (AP: + 0.8, ML: +/- 0.3, DV: −3.8) as depicted in **Figure 1A**. A total volume of 250 nL of hM4Di, hM3Dq, or mCherry was infused into the dSep using a 0.5 uL Hamilton Neuros syringe at a flow rate of 0.05 uL/min for 5 min. The syringe was then left in place during a 5 min diffusion period and retracted over 5 min as previously described [42]. Viral constructs obtained from Addgene for use in these experiments were AAV8-hSyn-hM4Di-mCherry (#50475), AAV8-hSyn-hM3Dq-mCherry (#50474), and AAV8-hSyn-mCherry (#114472). Post-operative care consisted of a single prophylactic treatment with buprenorphine (0.1 mg/kg) and 3 days of maintenance on a liquid acetaminophen solution during recovery.

**Figure 1:**
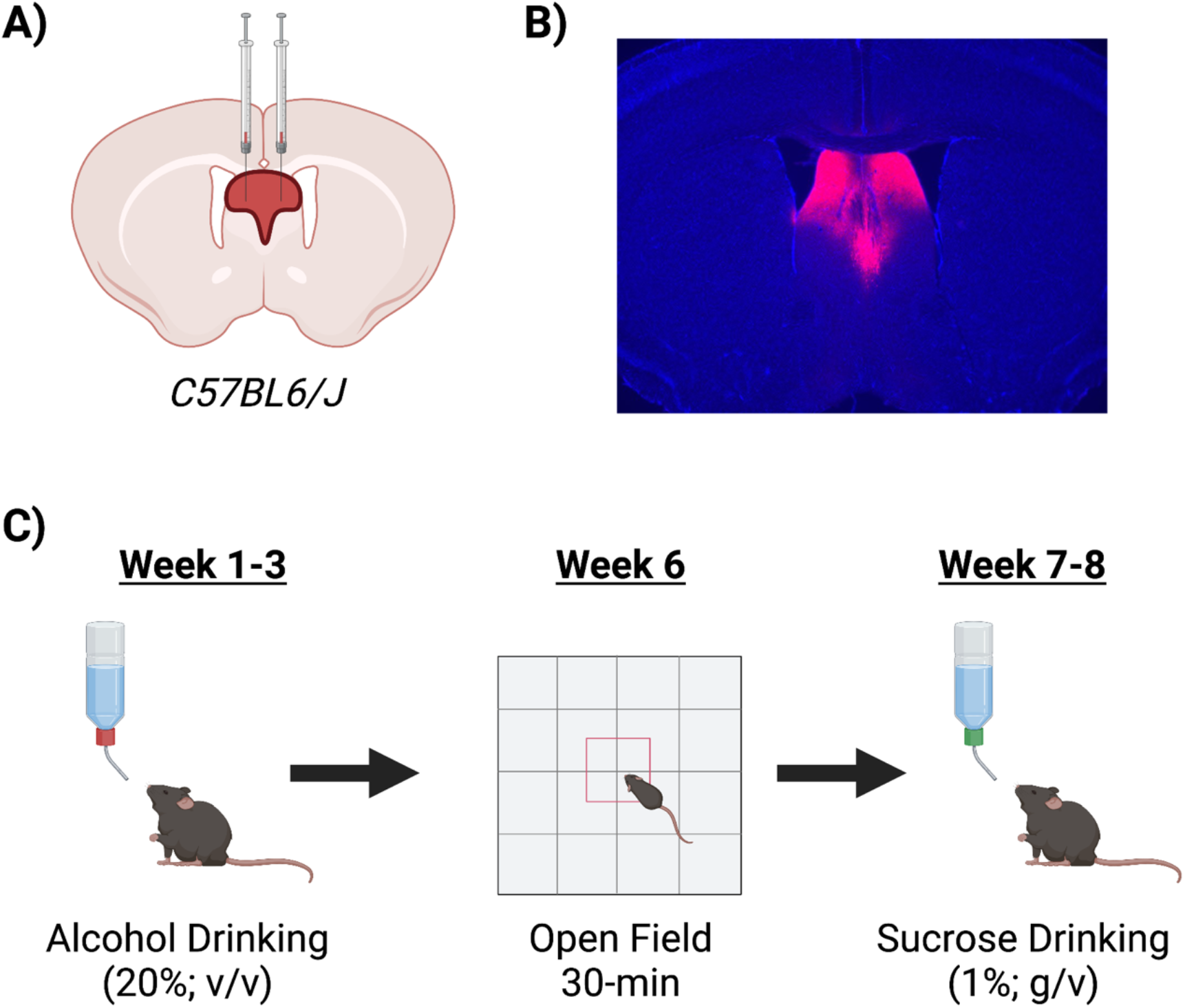
Viral Expression and Timeline of Experimental Procedures. **A)** AAVs harboring hM4Di, hM3Dq, or mCherry were infused into the dSep in male and female C57BL/6J mice 4 weeks prior to testing. **B)** Expression of mCherry was confined to the dorsal LS with some spillover into the dorsal edge of the MS. **C)** Experimental timeline depicting alcohol drinking, open field testing, and sucrose drinking.

### 2.3. Behavioral Testing

After surgery, mice were given 2 weeks to recover undisturbed in the home cage and then were habituated to intraperitoneal (ip.) injections of 0.9% saline (0.01 ml/kg) once daily for a week. Saline injections served as vehicle for all studies. After habituation to injections, mice began the alcohol drinking phase of testing, which occurred over 2-3 consecutive weeks. Following the final week of alcohol drinking, mice were left undisturbed in their home cage for 2 weeks. Mice were then tested in an open field arena to assess locomotor activity. The following week, mice began the sucrose drinking phase of the experiment for 2 consecutive weeks. A diagram depicting the timeline of experimental testing is shown in **Figure 1C**.

### 2.4. Drinking Procedures

Alcohol bottles were presented 3-hrs into the dark cycle during 2-hr limited-access drinking sessions for 5 consecutive days (Mon-Fri). Mice received daily saline injections 30-min prior to alcohol drinking. This modified drinking in the dark (DID) procedure was selected because it generates binge-like levels of alcohol drinking and BACs [43]. Most importantly, vehicle/drug testing can be conducted in a within-subjects fashion during a single test week, minimizing the potential for drinking differences or environmental disturbances between test weeks that is possible in the typical DID model. After habituation to alcohol drinking and saline injections (1-2 weeks), average weekly alcohol intake was calculated for each mouse and used to separate subjects evenly into drug treatment groups (vehicle vs CNO) in a balanced within-subjects design. During the Test week, all mice received vehicle injections prior to drinking on Day 1 and CNO testing occurred across Day 2-5 in a crossover design such that each subject received a single dose of CNO (3 mg/kg; HelloBio, HB6419). CNO was prepared daily in saline and delivered at 0.01 ml/kg volume 30-min prior to drinking.

Sucrose drinking was assessed in an identical fashion to alcohol drinking. Briefly, mice were given access to a single bottle containing sucrose (1%; w/v) for 2-hr a day, 3-hr into the dark cycle, Monday-Friday for two consecutive weeks. Testing occurred during the second week of drinking across Days 2-5 in a counterbalanced, crossover design.

### 2.5. Open Field Testing

Mice were left undisturbed in their home cage for 2 weeks following alcohol drinking and then activity chambers were used to assess locomotor activity. Subjects were tested in a SuperFlex open field arena (Omnitech Electronics, Accuscan, Columbus, OH) and proprietary Fusion v6.5 software was used to track beam breaks to determine total distance traveled during 30-min testing sessions. Briefly, the open field arena measured 60 cm wide × 60 cm long × 40 cm deep and a single light was positioned above the center of the apparatus (~500 lx). All mice were habituated to travel outside of the colony room for testing by placing cages on a covered cart and transporting to the behavioral testing room where they remained undisturbed for 8-hrs. The following day, mice were transported to the behavioral testing room, given a 2-hr acclimation period, then placed into the locomotor activity apparatus 30-min after vehicle injection. The goal of this experiment was to assess the effect of dSep inactivation/activation on locomotor behavior in a within-subjects design. Thus, a single session was used to habituate all mice to the locomotor apparatus prior to testing to control for novelty and order effects. Mice were handled in an identical fashion the following 2 days and tested with vehicle or CNO in a counterbalanced, crossover design. Distance traveled (cm) was measured in 1-min bins for 30 minutes during the two testing sessions and cumulative distance traveled was analyzed. Additionally, cumulative time spent in the center (10 cm) of the open field arena was collected to assess effects on anxiety-like behavior.

### 2.6. Immunohistochemistry

Upon completion of behavioral testing, all mice were deeply anaesthetized with Avertin (250 mg/kg; ip.) and transcardially perfused with ice cold saline (15 mL) followed by 4% paraformaldehyde (PFA; 15mL). Brains were extracted and post-fixed in PFA for 24-hrs. Tissue was sliced in 40 uM sections using a Vibratome and serial sections containing the entirety of the dSep were processed for immunohistochemistry to visualize the mCherry tag. Tissue was washed in 1xPBS, permeabilized in 0.5% Triton X-100 for 30-min, and blocked in 0.1% Triton X-100 and Normal Donkey Serum (NDS; 10%) for 1-hr. Tissue was incubated for 24-hr in the primary antibody solution that consisted of the blocking solution and mouse anti-RFP (1:500; Rockland Immunohistochemicals). The following day, tissue was washed in 1xPBS and incubated in a secondary solution that consisted of Alexa-Fluor 594 donkey anti-mouse (1:200; Jackson Immuno) in 0.1% Triton X-100 for 2-hr. Tissue was washed in 1xPBS, mounted on Permafrost Plus slides, and coverslipped for Vectashield Hardset with DAPI. Slides were imaged on a Keyence microscope and viral expression was verified within the dSep through visualization of mCherry within the target region. A depiction of the general viral footprint and spread within the dSep is shown in **Figure 1A** and a representative image of mCherry expression is shown in **Figure 1B**.

### 2.7. Statistical Analysis

The primary dependent variables for drinking experiments were alcohol intake (g/kg) and sucrose intake (mL/kg). Alcohol and sucrose drinking data after vehicle and CNO challenge were analyzed by ANOVA, with Sex and Virus (hM4Di; hM3Dq; mCherry) as between-subjects factors and Drug (Vehicle; CNO) as a repeated factor. Individual drinking values for vehicle and CNO testing days were then used to calculate the percent change in intake relative to vehicle ([(CNO-Vehicle) / (Vehicle)]*100). These data were then analyzed by ANOVA with Sex and Virus (hM4Di; hM3Dq; mCherry) as between-subjects factors and Drug as a repeated measure. For open field testing, cumulative locomotor activity and center time were analyzed by ANOVA with Sex and Virus as between-subjects factors and Drug as a repeated measure. Significant factor interactions were further evaluated with planned post hoc comparisons with Holm-Sidak corrections for effect of CNO treatment relative to vehicle within each experimental condition. Alpha was set to p< 0.05 for all analyses. A total of 8 mice were excluded from analysis due to unilateral AAV expression visualized by the mCherry tag within the dSep.

## 3. RESULTS

### 3.1. Chemogenetic Activation of the dSep Increases Alcohol Intake in Male Mice

The primary objective of this experiment was to determine the effect of chemogenetic manipulation of the dSep on binge-like alcohol consumption. Alcohol drinking data (g/kg) for male and female mice harboring hM4Di, hM3Dq, or mCherry in the dSep when challenged with vehicle and CNO is shown in **Fig. 2A**. Mice were treated with vehicle and CNO prior to a 2-hr limited access drinking session in a within-subjects, counterbalanced design. Consistent with the literature, a main effect of Sex *[F (1, 43) = 9.41, P= 0.004]* indicated that females generally consumed more alcohol than males in the DID model of binge drinking. A Sex*AAV interaction was observed *[F (2, 43) = 3.23, P= 0.049]* and the 3-way Sex*AAV*Drug interaction neared significance *[F (2, 43) = 2.58, P= 0.088]*. Planned comparisons post hoc analysis revealed that male hM3Dq-exressing mice consumed significantly more alcohol (+0.91 g/kg, +/- 0.33) when treated with CNO compared to vehicle (P= 0.008). This effect was not observed after CNO challenge in female mice harboring hM3Dq in the dSep (P= 0.57). Silencing (hM4di) of the dSep did not affect alcohol drinking in male and female mice nor did CNO influence drinking in mCherry control groups (Ps> 0.3).

**Figure 2:**
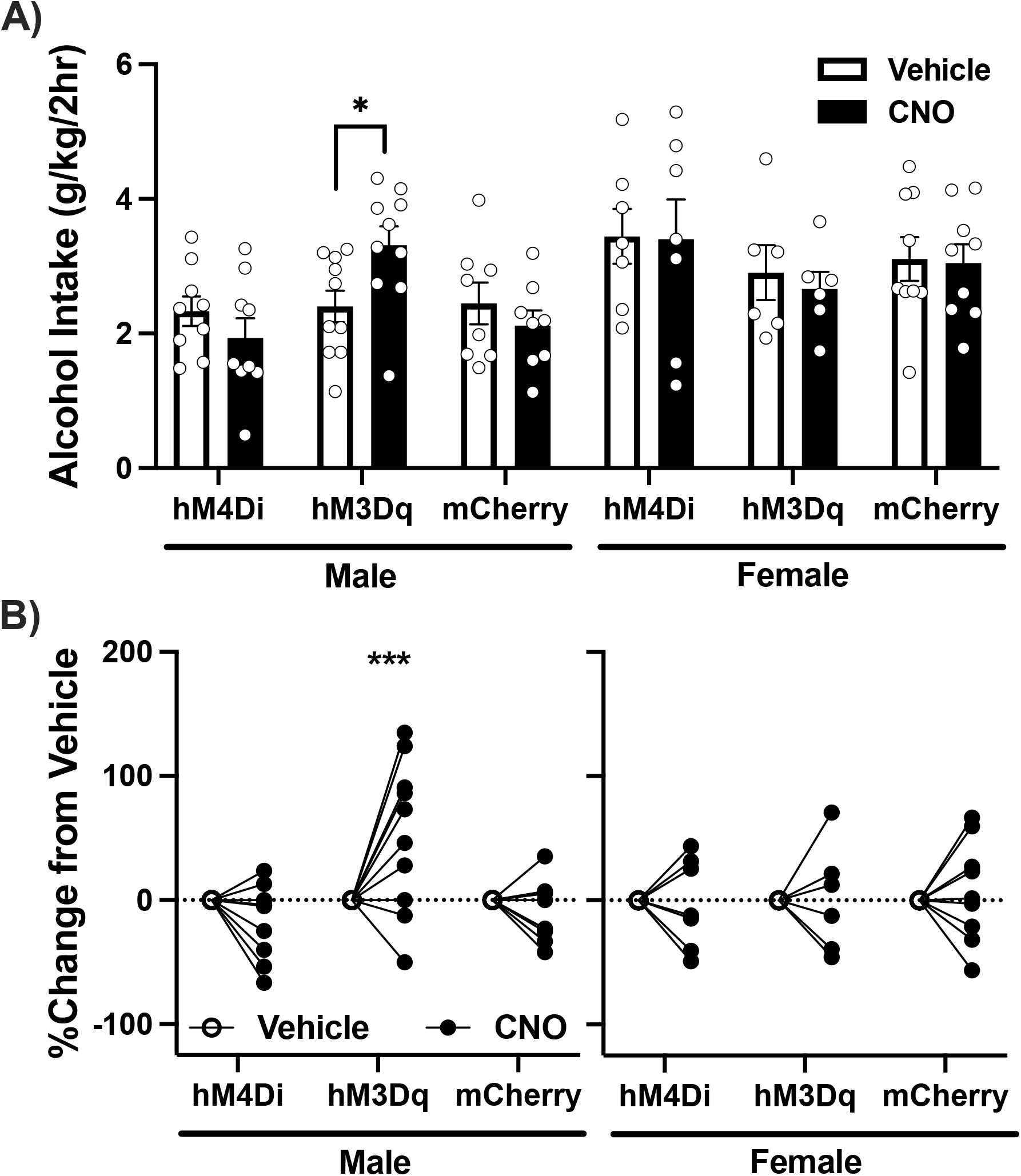
Chemogenetic Activation of the dSep Increases Alcohol Intake in Male Mice. **A)** Average alcohol intake (g/kg) after vehicle or CNO challenge. Alcohol drinking was greater in mice expressing hM3Dq in the dSep after CNO treatment when compared to vehicle (*P= 0.008). There was no effect of CNO treatment in the female hM3Dq group and CNO was without effect in hM4Di and mCherry conditions. **B)** Alcohol intake expressed as a percent change from vehicle after CNO challenge. Treatment with CNO in mice expressing hM3Dq within the dSep resulted in a significant increase in alcohol intake in males relative to vehicle (***P< 0.001). No significant change in intake was observed in hM4Di- or mCherry-expressing males. Lastly, there was no effect of experimental manipulation on the relative change in alcohol intake in female mice.

Drinking data were then analyzed as a percent change in alcohol intake for individual subjects relative to vehicle after CNO treatment (**Fig. 2B**). ANOVA revealed a near significant main effect of AAV *[F (1, 43) = 3.14, P= 0.053]* and a Sex*AAV*Drug interaction *[F (1, 43) = 3.26, P= 0.048]*. Post hoc analysis indicated that CNO treatment in hM3Dq-expressing mice resulted in an increase (+52.03%; +/- 13.3) in alcohol consumption in males (P< 0.001) but did not influence drinking in females (P= 0.95). No effect was observed in male hM4Di and mCherry groups, nor in female hM4Di, hM3Dq, and mCherry groups (Ps> 0.2). Together, these data suggest that activation of the dSep promoted alcohol drinking selectively in male mice.

### 3.2. The dSep Bidirectionally Modulates Locomotor Activity in Male Mice

Because the septal region contributes to locomotion [26,44], mice were tested in an open field arena to assess the effect of dSep manipulation on cumulative locomotor activity (**Fig. 3A**). After habituation to the locomotor boxes, mice were challenged with vehicle and CNO in a counterbalanced, within-subjects design. ANOVA revealed a main effect of AAV *[F (2, 43) = 5.35, P= 0.008]* and an AAV*Drug interaction *[F (2, 43) = 15.15, P< 0.001]*. A Sex*AAV*Drug interaction *[F (2, 43) = 6.02, P= 0.005]* was observed and post hoc analysis indicated that locomotor activity was affected in male hM4Di- and hM3Dq-expressing groups after CNO challenge. More specifically, CNO resulted in a decrease in locomotor activity (−1110.03 cm; +/- 470.26; P= 0.023) in male mice harboring hM4Di in the dSep, and this effect neared significance in females (−1010.61 cm; +/- 533.23; P= 0.065). Activation (hM3Dq) of the dSep after CNO treatment resulted in an increase in locomotor activity (+3164.28 cm; +/- 446.13) in males (P< 0.001) and was without effect in females (P= 0.63). Lastly, CNO had no effect on locomotor activity in male and female mice expressing mCherry in the dSep (Ps> 0.7). Time spent in the center of the open field was calculated and shown in **Supplemental Fig. 1A**. Briefly, female mice generally spent less time in the center of the open field supported by a main effect of Sex *[F (1, 43) = 4.58, P= 0.038]*. CNO did not affect center time *[F (1, 43) = 0.169, P= 0.683]* and post hoc indicated no differences in center time in any experimental condition when compared to CNO (Ps>0.2).

**Figure 3:**
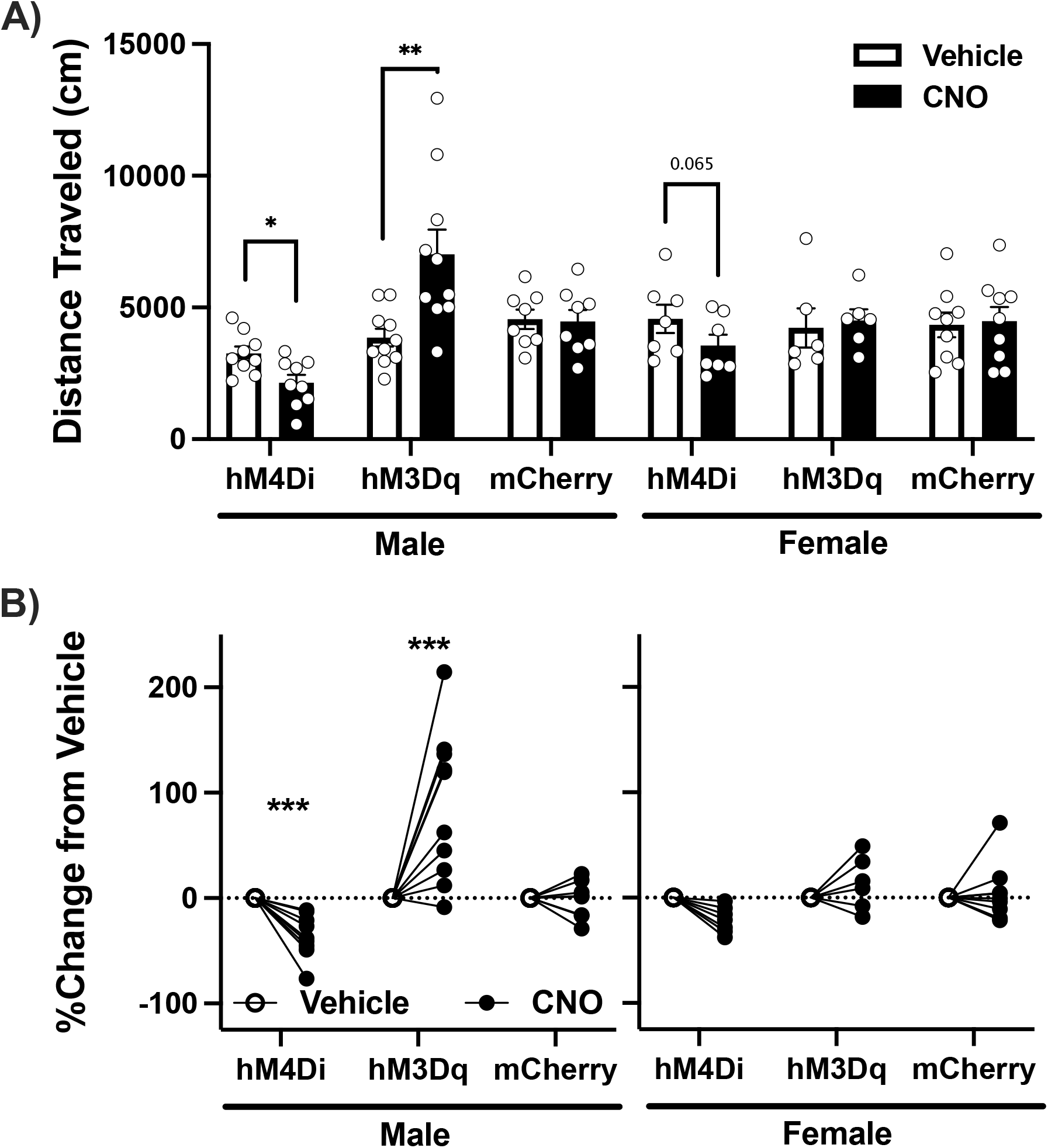
The dSep Bidirectionally Modulates Locomotor Activity in Male Mice. **A)** Cumulative distance traveled (cm) during an open field test. Compared to vehicle, CNO treatment resulted in lower distance traveled in male mice expressing hM4Di in the dSep compared to vehicle (*P= 0.023). Greater distance traveled was observed in the male hM3Dq group compared to vehicle (**P< 0.001). Female groups were not affected by CNO although there was a trend toward a decrease in distance traveled in the hM4Di condition (P= 0.065). **B)** Locomotor activity expressed as a percent change from vehicle after CNO challenge. Treatment with CNO in mice expressing hM4Di within the dSep resulted in a significant decrease in locomotor activity relative to vehicle in males (***P< 0.001). Conversely, activation of the dSep in males expressing hM3Dq resulted in increased locomotor activity (***P< 0.001). No significant change in locomotion was observed in female mice.

Analysis of locomotor activity when calculated as a percent change from vehicle indicated a similar pattern as cumulative distance traveled (**Fig. 3B**). There was a main effect of AAV *[F (2, 43) = 17.12, P< 0.001]* and a Sex*AAV*Drug interaction [F (2, 43) = 6.47, P= 0.003] indicating that locomotor activity was selectively impacted in male mice. Silencing (hM4Di) of the dSep decreased cumulative locomotor activity (−35.65%, +/- 12.5) in male mice (P= 0.007) but did not affect distance traveled in females (P= 0.14). In contrast, activation of the dSep increased locomotor activity (+87.13%, +/- 11.86) in males but was without effect in females (P= 0.38). Together, these data indicate a bidirectional effect of dSep manipulation selectively in male mice.

### 3.3. Activation of the dSep Promotes Sucrose Drinking in Male and Female Mice

A final experiment was conducted to determine the effect of dSep manipulation on sucrose drinking (**Fig. 4A**).This experiment was conducted in a similar fashion to alcohol drinking, with vehicle and CNO being tested in a within-subjects, counterbalanced fashion. A main effect of Sex *[F (1, 43) = 99.89, P< 0.001]* indicated that females consumed significantly more sucrose than males in the 2-hr limited access drinking session. ANOVA also revealed an AAV*Drug interaction *[F (2, 43) = 4.19, P= 0.022]* and post hoc analysis supported an effect of dSep activation to increase sucrose drinking. Activation of the dSep promoted sucrose drinking (+6.59 mL/kg; +/- 3.59) in male mice (P= 0.027) and a similar trend toward an increase in intake (+10.62 mL/kg; +/- 4.63) was observed in females (P= 0.073). Silencing of the dSep did not affect sucrose drinking in male or female mice and CNO was without effect in mCherry conditions (Ps> 0.5).

**Figure 4:**
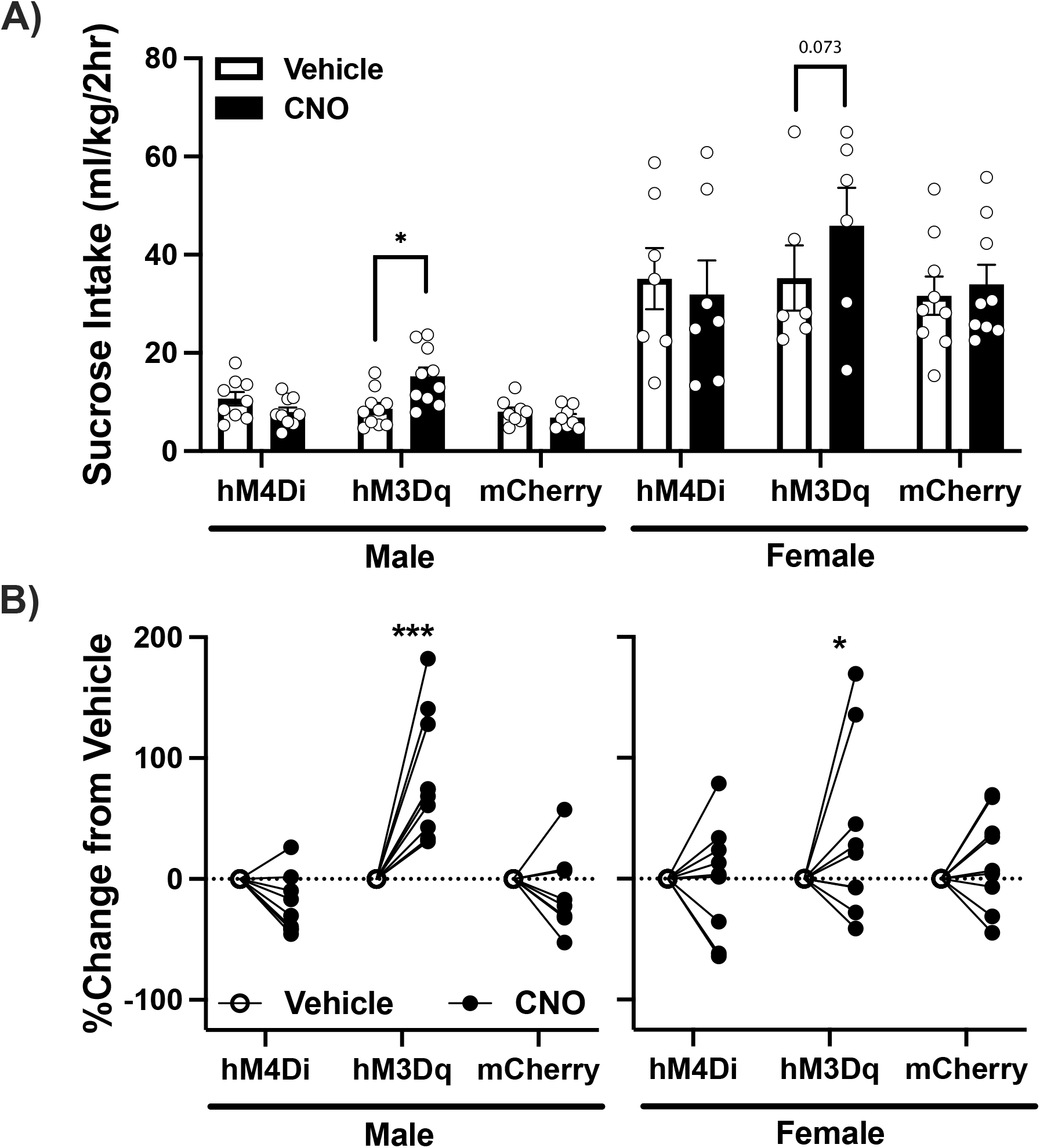
Activation of the dSep Promotes Sucrose Drinking in Male and Female Mice. **A)** Sucrose intake (ml/kg) after vehicle or CNO challenge. Sucrose drinking was significantly greater after treatment with CNO in male mice expressing hM3Dq within the dSep (*P= 0.008) and this effect neared significance in females (P= 0.073). There was no effect of CNO in hM4Di and mCherry conditions. **B)** Sucrose intake expressed as a percent change from vehicle after CNO challenge. Treatment with CNO in mice expressing hM3Dq within the dSep resulted in a significant increase in sucrose consumption relative to vehicle in males (***P< 0.001) and females (*P= 0.021). There was no change in sucrose intake in hM4Di or mCherry conditions.

Analysis of sucrose drinking data when expressed as a percent change in intake relative to vehicle support the effect of dSep activation to increase sucrose intake (**Fig. 4B**). ANOVA revealed an AAV*Drug interaction *[F (2, 43) = 10.80, P< 0.001]* and further post hoc analysis support a general increase in sucrose drinking in male and female mice. More specifically, dSep activation resulted in an 83.70% +/- 15.36 increase in sucrose drinking in males (P< 0.001) and an 47.59% +/- 19.83 increase in females (P= 0.021). Silencing of the dSep did not alter the relative change in sucrose drinking (Ps<0.2) nor did CNO alter intake in mCherry control groups (Ps>0.4).

## 4. DISCUSSION

The septal complex has been implicated in reward and various aspect of drug addiction, yet no studies have interrogated this structure in the context of binge-like alcohol consumption. Here, we report that chemogenetic activation (hM3Dq) of the dSep increased voluntary alcohol drinking in male mice in a modified-DID model. Alcohol intake was not affected in females, yet activation of the dSep resulted in increased sucrose drinking in both males and females suggesting a general role for the region in consummatory behavior. Silencing (hM4Di) of the dSep was without effect on alcohol or sucrose intake independent of sex. Lastly, activation or inhibition of the dSep increased/decreased locomotor activity respectively in males, but did not influence distance traveled in females. Therefore, the general increase in alcohol and sucrose intake in male mice may be due to a nonspecific increase in locomotor activity. Together, these data support a role for the dSep in promoting voluntary consumption of a palatable reward and suggests a role for the structure in alcohol drinking and locomotion in a sex-dependent fashion.

The septum is a centrally located subcortical limbic structure and is comprised of two distinct subregions: the lateral septum (LS) and medial septum (MS) [45]. The LS is situated in the dorsolateral region of the septal complex and is thought to promote goal directed behavior and learning by relaying spatial information from the hippocampus to the ventral tegmental area (VTA) for reward processing [46,47]. The present studies were planned to selectively target the LS yet rigorous histological review revealed an unexpected extension of mCherry expression into the dorsal edge of the MS. The MS is most well-known for its cholinergic contribution to septohippocmpal processing and involvement in the regulation of theta oscillations [48,49]. Breese and colleagues demonstrated MS responsivity to alcohol and a role for the structure in alcohol-induced sedation and seizure activity [36,50–52]. However, selective MS ablation resulted in increased water intake whereas LS ablation was without effect [53]. Stimulation of the MS conversely decreased drinking [54] suggesting bidirectional MS control over water intake that is in the opposite direction of the effects observed in the present studies. Thus, it is unlikely that the increase in alcohol drinking and sucrose intake after dSep activation is driven by the MS but we cannot attribute observed effects in our experiments to either the LS or MS alone. Discrete differentiation of the LS and MS is important for further study and identification of cell-types that promote drinking behavior will be of interest to the field.

Mechanistic study of the septal contribution to drug-seeking has been largely limited to cocaine and opioids [40,41,55,56]. However, there is ample evidence to suggest that the septal complex is responsive to alcohol. Alcohol generally suppresses c-fos activity within the septum and decreases catecholamine levels after both acute and chronic treatment [34,35,57,58]. However, findings are inconsistent in that intracerebroventricular alcohol infusion increased c-fos expression within the LS and promoted the development of a conditioned taste preference for a saccharin solution [59]. These findings suggest that alcohol promotes activity within the septum that is involved in reward processing and there is strong evidence supporting LS regulatory control over mesolimbic function. Indeed, the septum projects to both the VTA and NAc and these circuits have been hypothesized to promote reward-related behaviors [60,61], including excessive drinking behavior. Because pharmacological inactivation of the LS blocks the ability of alcohol to elevate extracellular dopamine within the NAc [62], it is reasonable to suspect that activation of the dSep in the present experiments could promote alcohol (and sucrose) intake through activation of canonical mesolimbic reward pathways. Furthermore, c-fos activity is elevated within the septum during withdrawal from alcohol suggesting that neuroadaptations occur within this structure over time [38]. The excessive alcohol drinking behavior observed after dSep activation is similar to the escalation of intake observed in dependence models and suggests a shared neurobiological mechanism. Another possible projection site of the dSep that modulates appetitive behavior is the hypothalamus. Both the LS and MS send projections to hypothalamic subregions and LS projections to the arcuate nucleus and lateral hypothalamus (LH) have been implicated in feeding [63] behavior and reinstatement of cocaine-seeking [40], respectively. The LH is of particular interest given that activation of GABAergic neurons therein promotes alcohol drinking and general consummatory behavior [64,65]. Thus, the VTA and LH serve as two likely downstream septal projection sites mediating alcohol drinking behavior.

The septal complex is involved in locomotion and regulation of speed, velocity, and acceleration [66,67]. Our findings support previous work and indicates that the dSep exerts bidirectional control over locomotor activity in males. The septum, LS specifically, is hypothesized to drive a movement-value signal that is sent downstream to promote goal directed behavior [26]. In fact, hippocampal place-like cells within the LS are highly active in reward paired contexts [67] and manipulation can result in altered frequency and preservation of learned behaviors [68,69]. Thus, it is not surprising to see changes in locomotion in the open field task and this general increase in activity may present as increased consummatory behavior in the drinking tasks. Beyond this, if the septum is involved in reward learning and integration of movement involved in the behavior, dSep activation may promote drinking behavior by activating circuitry that enhances the valence of the rewarding stimulus, presenting as increased alcohol or sucrose drinking behavior.

Perhaps the most intriguing finding of the present study is that of sex differences in the outcome of dSep activation on alcohol and sucrose drinking behavior. Females generally consume more alcohol and sucrose than males in the DID model and there is much interest in identifying sexually dimorphic regions that promote this behavior [70–73]. Sex differences have been noted within the septum [74–76] and the ability of dSep activation to promote alcohol drinking in males is of particular interest. Here we report that dSep activation elevated alcohol intake in males (+52%) to a level that matched intake in females. However, sucrose drinking was equally affected in males and females. It is possible that a ceiling effect was reached for alcohol consumption in females obscuring the potential to observe a hM3Dq-mediated increase in intake, and this possibility makes interpretation difficult. However, females consumed 3.4 g/kg in 2-hrs and we have previously shown upwards of 4 g/kg intake in C57BL/6J mice after pharmacological intervention [70], suggesting room to observe an increase in alcohol drinking. Lastly, it is surprising that dSep manipulation was without effect on locomotor activity in females. We have previously found greater distance traveled in female C57BL6/J mice under similar conditions in an open field arena [70] but no differences were observed in the present study. In contrast to previous work, mice were thoroughly habituated to the open field to allow for within-subjects testing and this may obscure heightened locomotion in females. Anxiety-like behavior, as determined through time spent in the center of the open field, remained unaffected by dSep manipulation. These data suggest that dSep activation promoted locomotor activity, but is not innately anxiogenic as others have speculated. Clearly, this single observation in the open field task is not conclusive, and habituation to the apparatus can confound this measure. However, the dSep likely functions to promote learned locomotor action linked to a specific context, and more complex assays for anxiety-like behavior linking the context to a specific outcome may be more appropriate for further septal interrogation.

The septum remains relatively understudied within modern alcohol research amidst a plethora of data supporting a role for the structure in behaviors and processes involved in the regulation of drinking behavior. Novel techniques within the preclinical toolkit allow for discrete manipulation of neuroanatomic targets such as the septal complex and the present findings indicate that the dSep plays a faciliatory role in promoting alcohol and sucrose drinking in sex-dependent fashion. However, much work is necessary to determine the exact mechanism that facilitates these effects. Because activation of the dSep increased alcohol drinking, an intriguing future direction is to determine if escalated alcohol intake in dependence models is dependent upon dSep activity. Lastly, discrete manipulation of the LS and MS and select cell-types within will be important for future studies.

---

**Supplemental Figure 1.**
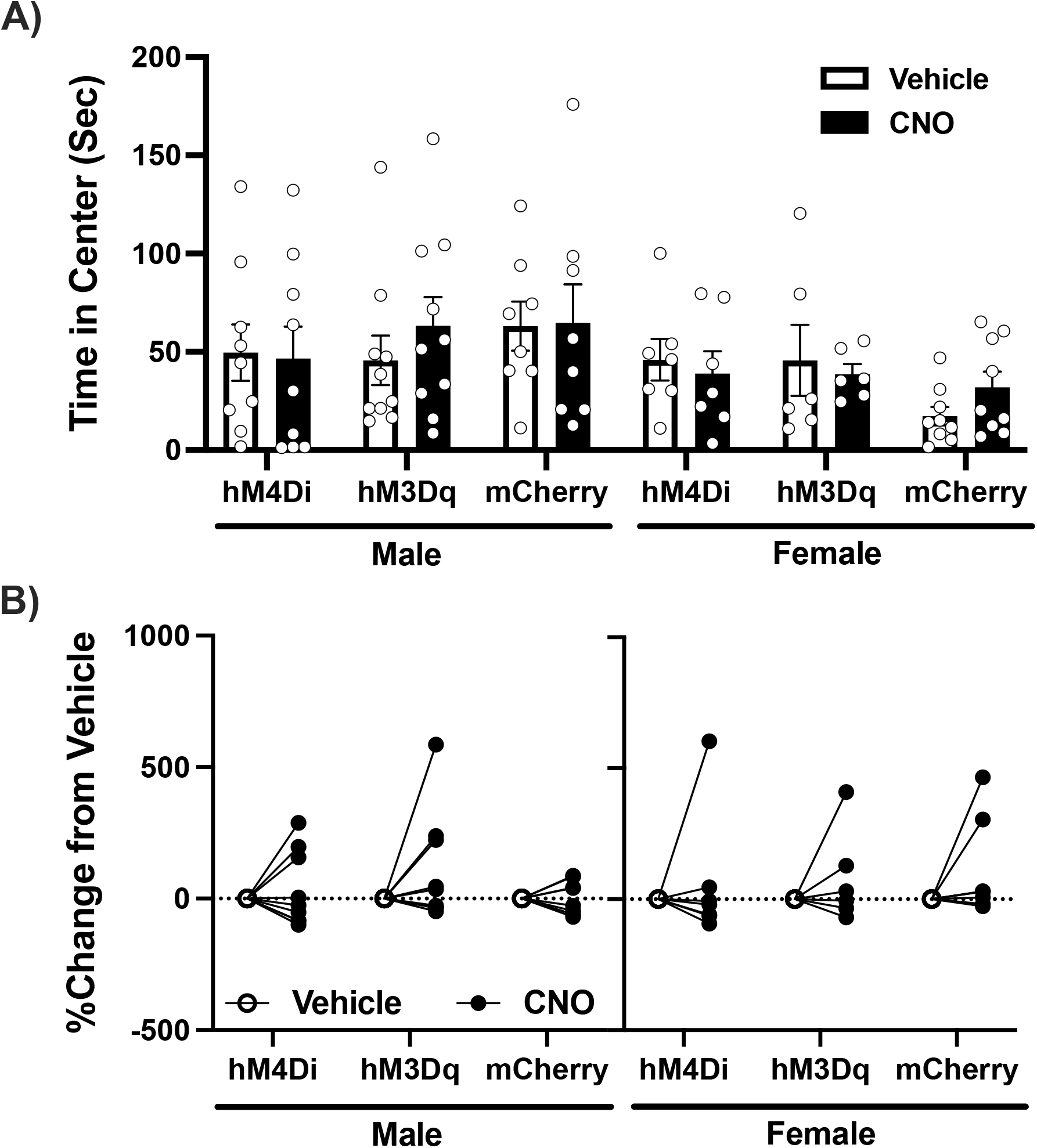
Manipulation of the dSep Does Not Affect Anxiety-Like Behavior in an Open Field Arena. **A)** Cumulative spent in the center of the open field. No differences were detected in any condition after CNO challenge. However, females generally exhibited less center time than males (P= 0.038). **B)** Time spent in center expressed as a change relative to vehicle. CNO treatment did not significantly alter center time in any experimental condition.

## 5. ACKNOWLEDGMENTS

Graphical abstract was created with BioRender.

